# Discovery of latent clinical phenotypes through optimal transport of brain dynamics

**DOI:** 10.64898/2026.11.13.766403

**Authors:** Luca Taffarello, Giacomo Barzon, Benedetta Mariani, Federico Calesella, Antonio Gallo, Manuela Altieri, Alessandro Pasquale De Rosa, Alessandro D’Ambrosio, Fabrizio Esposito, Stefano De Marchi, Antonino Vallesi, Michele Allegra, Alessandra Bertoldo, Samir Suweis, Marco Zorzi

## Abstract

The increasing availability of large clinical neuroimaging datasets has enabled the discovery of objective, data-driven clinical phenotypes. However, current approaches typically rely on structural or static functional metrics, thereby overlooking the rich information embedded in brain dynamics. To address this limitation, we developed a computational framework combining a Gaussian Hidden Markov Model (HMM) with Optimal Transport (OT) theory. By representing individual brain dynamics within a shared latent state space, our approach quantifies how much one subject’s state-transition dynamics would need to be modified to match another subject’s state occupancy distribution, providing a pairwise measure of inter-individual variability that captures temporal aspects of brain function. We applied this framework to multiple sclerosis (MS), a condition where substantial within-group heterogeneity challenges traditional clinical classification. Analyzing resting-state functional magnetic resonance imaging data from 122 patients with relapsing-remitting MS and 97 matched healthy controls, the HMM identified three latent states with distinct connectivity patterns.

While patients showed an altered temporal organization of recurrent brain states relative to healthy controls, reflected in selective differences in state occupancy and dwell time, we focused on quantifying heterogeneity specifically within the MS cohort. Clustering patients according to OT-based dissimilarities revealed two subgroups with distinct clinical disability and structural disease burden. Crucially, this stratification was not detectable when applying clustering procedures to static functional connectivity, supporting the added value of dynamical features. Our results establish a general methodological framework for extracting clinically relevant phenotypes from temporal brain dynamics, with potential applications across neurological and psychiatric conditions characterized by phenotypic heterogeneity.

**Author summary:** Even at rest, the brain continually shifts among recurring patterns of coordinated activity across regions, commonly called brain states. Brain imaging studies aimed at characterizing individual differences often focus on average relationships between regions, potentially overlooking how activity evolves over time. We developed a computational framework that identifies these states and quantifies differences in their dynamics across individuals. Specifically, we estimate how much one person’s transitions between states would need to change for their brain to spend the same proportion of time in each state as another person’s. We applied this framework to multiple sclerosis, where disability and brain damage vary widely among patients. Using brain scans acquired at rest, we found that the distribution of time across brain states differed between patients and healthy controls. More importantly, without using clinical information, our framework identified two groups of patients who differed in disability, brain atrophy, and lesion burden, whereas grouping patients by average connectivity did not reveal comparable clinical distinctions. Our findings suggest that quantifying differences in brain-state dynamics across individuals can reveal clinically relevant information that averaged measures may miss. With further validation, this framework could help characterize clinical heterogeneity in multiple sclerosis and other neurological conditions.

## Introduction

Traditionally, the clinical phenotyping of neurological and psychiatric diseases relied heavily on behavioral markers. The advent of large-scale neuroimaging has since enabled the delineation of phenotype boundaries based on anatomical and functional correlates of behavioral or cognitive impairment [1, 2]. While this paradigm has yielded significant insights into psychiatric [3, 4], neurodegenerative [5–7], and neurodevelopmental diseases [8], it typically relies on either structural markers or static, time-averaged functional connectivity (FC). By focusing on static features, these approaches overlook the rich information embedded in dynamic activity patterns, which are known to be essential for explaining variance in behavioral traits and higher cognition [9–13]. Despite the documented alterations in brain dynamics across various conditions [14–16], the use of whole-brain dynamical markers for clinical stratification remains relatively underexplored [17].

Here, we propose a principled framework to characterize inter-individual variability in whole-brain dynamics for subject stratification. Unlike clustering methods based on engineered features, our approach provides a geometrically grounded dissimilarity measure rooted in optimal transport theory. We first employ a Gaussian Hidden Markov Model (HMM) [18, 19] to identify a set of latent states common to all subjects. Compared to sliding-window functional connectivity [14, 20, 21], HMMs offer more robust inference with less dependence on free parameters [22] and naturally represent dynamics as Markov transitions between discrete states, yielding probability distributions suitable for transport-theoretic analysis. While latent states are identified at the group level, individual dynamics are characterized by subject-specific state occupancy distributions and state-transition statistics. For each pair of subjects, we use the *Schrödinger bridge* formulation [23] of optimal transport (OT) to determine the *transport cost*, which measures the minimum modulation of the source subject’s state-transition dynamics needed to match the target subject’s latent-state occupancy distribution. This probabilistic formulation provides a stochastic-control interpretation of inter-individual differences in brain dynamics [24] and situates our approach within the emerging field of control-theoretic neuroscience [25]. While previous studies applied the Schrödinger bridge technique to estimate the transition cost between resting and task-evoked brain activity distributions [26, 27], we extend this approach to quantify inter-individual transport costs and construct a pairwise dissimilarity matrix. This enables fully unsupervised clustering and the embedding of subjects within a low-dimensional dynamic feature space.

To demonstrate the clinical relevance of this framework, we applied it to a large dataset of patients with relapsing-remitting multiple sclerosis (RRMS) and healthy controls (HCs) (Fig 1). Multiple sclerosis (MS) is a chronic inflammatory, demyelinating, and degenerative disease of the central nervous system [28]. While MS is traditionally classified into three phenotypes based on symptomatology (relapsing-remitting, primary progressive, and secondary progressive), significant within-group heterogeneity highlights the need to identify biologically grounded phenotypes [29–35]. Whereas previous dynamic connectivity studies in MS identified reliable patient-control differences and clinically relevant alterations associated with cognitive impairment, disability, and disease progression [36–40], they did not use brain dynamics for data-driven patient stratification. Using our optimal transport framework, we identified two dynamically distinct subgroups within the RRMS cohort that differed significantly in clinical disability and structural pathology, a stratification not recovered by static functional connectivity analysis.

**Fig 1.**
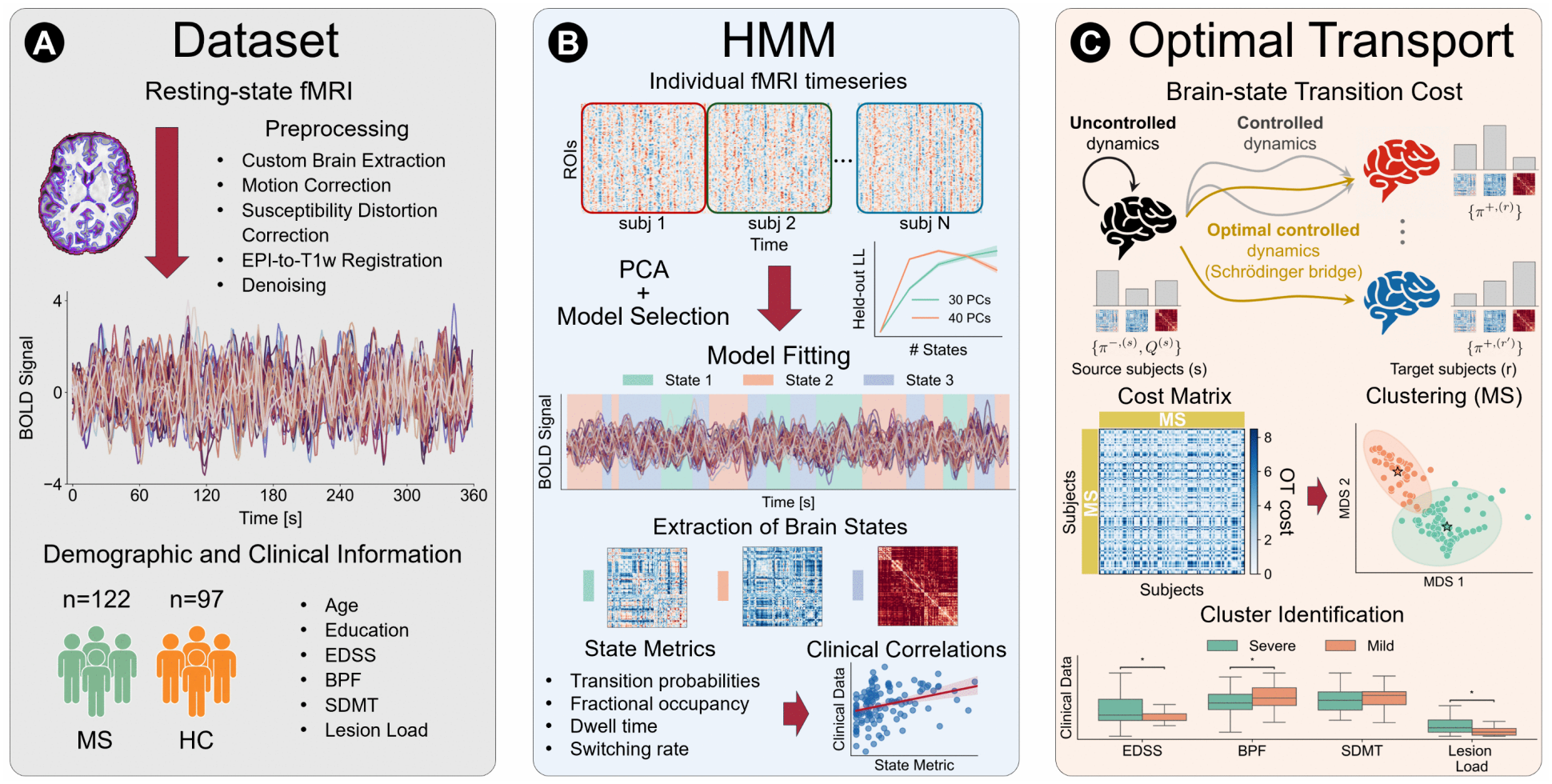
Workflow. (A) *Dataset and preprocessing*. Resting-state fMRI data were collected from 122 patients with multiple sclerosis (MS) and 97 healthy controls (HCs). Preprocessing was performed using fMRIPrep and XCP-D and included custom brain extraction, motion correction, BOLD-to-T1w co-registration, denoising, and extraction of parcel-wise time series using the Schaefer-100 atlas. Demographic information, clinical measures including the Expanded Disability Status Scale (EDSS) and Symbol Digit Modalities Test (SDMT), and MRI-derived measures including brain parenchymal fraction (BPF) and lesion load were also obtained. (B) *Hidden Markov Model (HMM) inference*. ROI-wise BOLD time series were standardized within each subject and dimensionality was reduced via principal component analysis (PCA). A Gaussian HMM was then fitted at the group level to identify recurrent brain states characterized by distinct covariance patterns of brain activity. Subject-specific state time courses were used to compute dynamic metrics including fractional occupancy, dwell time, and switching rate, which were subsequently related to clinical and imaging variables. (C) *Optimal transport (OT) analysis*. For each subject, posterior HMM estimates were used to characterize state occupancy distributions and transition dynamics. A Schrödinger bridge framework was used to estimate pairwise directional transition costs between subjects, quantifying the dissimilarity in their brain-state dynamics. The resulting cost matrix was symmetrized and used for clustering within the MS cohort. Clinical and imaging variables were subsequently compared across the identified subgroups to characterize their differences.

## Materials and methods

### Dataset

Magnetic resonance imaging (MRI) and clinical data were obtained from 122 patients with RRMS and 97 age- and sex-matched HCs recruited at the University of Campania Luigi Vanvitelli (Naples, Italy) [35]. All patients underwent neurological and neuropsychological evaluation, including the Expanded Disability Status Scale (EDSS) and the Symbol Digit Modalities Test (SDMT). MRI scans were acquired on a 3T GE Signa HDxt scanner (GE Healthcare, Milwaukee, USA) equipped with an eight-channel head coil. The imaging protocol included high-resolution 3D T1-weighted images (voxel size of 1 *×* 1 *×* 1.2 mm^3^) and 2D fluid-attenuated inversion recovery (FLAIR) images (voxel size of 0.47 *×* 0.47 *×* 3 mm^3^). Resting-state functional MRI (fMRI) data were acquired using an echo-planar imaging sequence (TR = 1500 ms; voxel size of 4 *×*4 *×*4 mm^3^; 240 volumes). Participants were instructed to remain still, awake, and with eyes closed during the fMRI acquisition. Brain tissue and lesion volumes were estimated from T1-weighted and co-registered FLAIR images using sequence adaptive multimodal segmentation (SAMSEG; FreeSurfer v7.1.1). Lesion load was quantified as total lesion volume, and brain parenchymal fraction (BPF) was calculated as the ratio of brain parenchymal volume to intracranial volume. Further details on participant recruitment, inclusion criteria, demographic and clinical characteristics, MRI acquisition, and the estimation of these structural measures are reported in [35].

### Preprocessing of MRI data

Functional and anatomical MRI data were preprocessed using fMRIPrep 24.1.1 [41, 42]. A custom anatomical module was incorporated to generate a high-quality, template-informed brain mask. T1-weighted images were first corrected for intensity inhomogeneities, then linearly and nonlinearly aligned to the MNI152 template. The resulting transformations were inverted to project a standardized template brain mask into native space, yielding a robust subject-specific mask. T1-weighted structural images also underwent surface reconstruction and tissue segmentation with FreeSurfer. Spatial normalization to MNI152NLin6Asym and MNI152NLin2009cAsym templates was performed using nonlinear ANTs registration. Functional preprocessing included creation of a reference volume, motion correction with MCFLIRT, and boundary-based co-registration of blood oxygen level-dependent (BOLD) data to the T1-weighted reference. Several confound time series and quality-control metrics were computed, including framewise displacement (FD), DVARS (the root mean square of voxelwise BOLD signal differences between consecutive volumes), global signals, and both anatomical and temporal CompCor components. Surface and grayordinate representations were generated by resampling BOLD data into fsLR space. Post-processing of fMRIPrep output was performed using the eXtensible Connectivity Pipeline-DCAN (XCP-D) [43]. BOLD data were despiked with AFNI’s 3dDespike. Denoising followed the aCompCor strategy, incorporating the top five white matter and cerebrospinal fluid components, 12 motion parameters (6 motion parameters and their derivatives), and high-pass cosine regressors. High-motion volumes (FD *>* 0.5 mm) were interpolated prior to band-pass filtering (0.01–0.1 Hz). The cleaned BOLD data were smoothed using a Gaussian kernel with a 6-mm full width at half maximum. Parcel-wise time series were then extracted using the 100-parcel version of the Schaefer atlas.

### Hidden Markov model

To characterize the temporal dynamics of large-scale brain activity, we employed a Gaussian HMM [18, 44]. This probabilistic framework models fMRI time series as transitions between a finite set of latent brain states. The HMM simultaneously estimates these latent states, each characterized by a multivariate Gaussian distribution of regional activity, along with the probabilities governing their temporal transitions. More explicitly, let *x*_*t*_ *∈* ℝ^*D*^ denote the vector of BOLD signals at time *t* from *D* regions of interest (ROIs), and let *z*_*t*_ *∈ {* 1, …, *K}*indicate the latent state active at that time. The temporal evolution of the latent states follows a first-order Markov process:

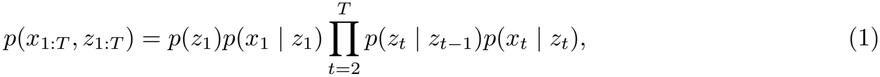

with 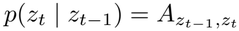 denoting the transition probabilities between states. The observation model for each state *k* is given by

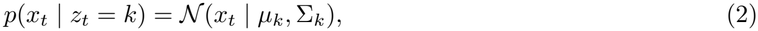

where *µ*_*k*_ andΣ_*k*_ represent the mean activation and covariance associated with state *k*. In the primary analysis, the state means were constrained to zero (*µ*_*k*_ = 0 for all *k*); results obtained with unconstrained state means are reported in Fig I in S1 Supporting Information.

Although the HMM can be estimated separately for each participant, the limited length of typical resting-state fMRI time series provides insufficient samples to reliably estimate all model parameters. Therefore, following common practice in the literature [45–49], a single group-level HMM was estimated jointly across patients and controls, providing a common state space for direct comparison between groups. Each subject’s ROI-wise time series was z-scored across time (zero mean and unit variance per ROI), and the standardized observations were concatenated across subjects for principal component analysis (PCA).

Dimensionality reduction was performed by retaining the first *M* components, thereby reducing the number of free parameters and improving the stability of covariance estimation and the statistical efficiency of model inference [18]. We selected *M* = 30, which explained approximately 80% of the total variance. The resulting *M* -dimensional PCA scores were used to fit the group-level HMM, treating each participant’s projected time series as an independent sequence so that no state transitions were modeled across subject boundaries. A common set of HMM parameters was estimated across subjects, while the state time courses were inferred separately for each participant. The model was estimated using the expectation-maximization algorithm implemented in the Dynamax library [50]. During the E-step, the algorithm computes the posterior probabilities of the latent states given the current parameters, and during the M-step it updates the transition matrix and the state-specific covariance matricesΣ_*k*_. Model fit was quantified using the marginal data log-likelihood, which under this independent-sequence formulation is additive across subjects:

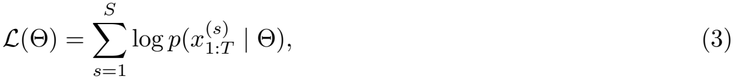

where Θ denotes the shared HMM parameters and *S* the number of subjects.

The number of hidden states (*K*) was selected using five-fold subject-wise cross-validation, stratified by diagnostic group. Within each fold, PCA was fitted exclusively on the training subjects and subsequently applied to the held-out subjects. For each fold, the HMM was fitted to the training sequences from 10 random initializations, and the solution with the highest training marginal log-likelihood was evaluated on the held-out subjects using the mean marginal log-likelihood. Cross-validation was performed for *K ∈ {*2, 3, 4, 5, 6, 7, 8} and repeated across *M ∈ {*25, 30, 35, 40, 45, 50} to assess the robustness of model-order selection to PCA dimensionality (see Fig A in S1 Supporting Information). Across PCA dimensionalities, held-out log-likelihood showed a marked increase from *K* = 2 to *K* = 3, followed by substantially smaller gains for additional states. This elbow-like pattern was consistent across *M*, and differences among high-*K* solutions were small relative to cross-validation variability. Based on this pattern and for the sake of model parsimony, we selected *K* = 3 for the primary analysis. The final HMM was fitted to the complete dataset using 20 random initializations, retaining the solution with the highest marginal log-likelihood. From the estimated model, we derived subject-specific metrics describing the temporal organization of brain-state activity: fractional occupancy (FO), mean dwell time (DT), and switching rate (SR). The FO of each state quantifies its posterior occupancy over time and was defined as

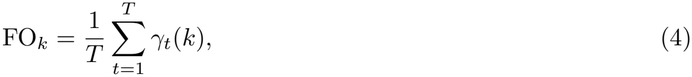

where *γ*_*t*_(*k*) = *p*(*z*_*t*_ = *k* |*x*_1:*T*_) denotes the posterior probability of state *k* being active at time *t*. For the computation of DT and SR, the most likely state sequence was obtained using the Viterbi algorithm [51]:

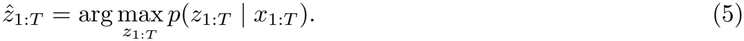

The DT of state *k* was defined as the average duration of time intervals during which the state remained active:

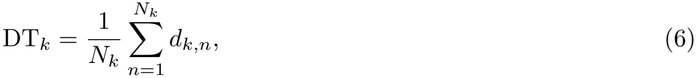

where *d*_*k,n*_ denotes the duration of the *n*-th interval in state *k*, and *N*_*k*_ is the total number of such intervals. The SR quantified the proportion of consecutive time-point pairs exhibiting a transition between distinct states:

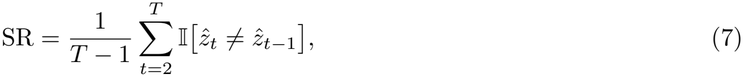

where I[*·*] is the indicator function.

### Transition cost between brain-state configurations via Schrödinger bridge

To quantify the dissimilarity between subjects in terms of their HMM-derived brain dynamics, we computed a transition cost between their respective brain-state configurations using the Schrödinger bridge framework [26, 27, 52]. For each subject *s*, we derived the joint probability matrix of consecutive latent states by averaging the posterior joint probabilities conditioned on that subject’s observations under the shared HMM:

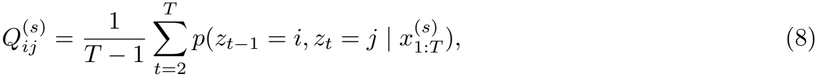

With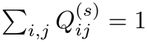. Thus, *Q*^(*s*)^ captures subject-specific transition statistics while accounting for uncertainty in latent-state assignments. These matrices were used as reference joint transition distributions for each subject’s resting-state dynamics. The row and column marginals of *Q*^(*s*)^ define the distributions of latent states before and after a transition, respectively:

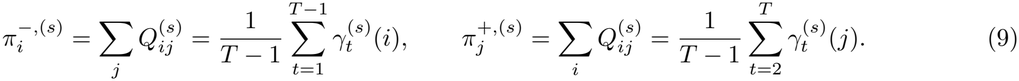

These marginals represent the average posterior state occupancy over all but the final or initial time point, respectively, and therefore converge to the fractional occupancy as the length of the time series increases. The Schrödinger bridge formalism provides a probabilistic approach to model how the latent-state occupancy distribution of a source subject *π*^*−*,(*s*)^ can be transformed into that of a target subject *π*^+,(*r*)^, using *Q*^(*s*)^ as the reference joint transition distribution. More specifically, it identifies a joint distribution *P* of consecutive latent states whose row and column marginals match *π*^*−*,(*s*)^ and *π*^+,(*r*)^, respectively. Among all such admissible joint distributions, the Schrödinger bridge solution is the one that minimizes the Kullback–Leibler divergence from *Q*^(*s*)^. In other words, this optimal *P* represents the controlled stochastic evolution connecting *π*^*−*,(*s*)^ to *π*^+,(*r*)^ while remaining as close as possible to the source subject’s reference (uncontrolled) dynamics encoded by *Q*^(*s*)^. We therefore defined the directional transition cost as

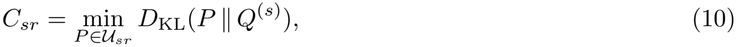

where 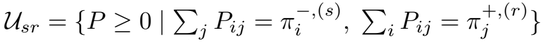 denotes the set of admissible joint transition probabilities (couplings) whose marginals match the source and target occupancy distributions. This formulation is equivalent to an entropy-regularized optimal transport problem [53, 54]:

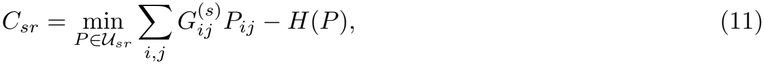

where 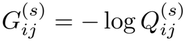 defines the transport cost matrix over latent states, derived from the reference joint transition distribution, and *H*(*P*) =*−* Σ_*i,j*_ *P*_*ij*_ log *P*_*ij*_ denotes the entropy term. The equivalent optimal transport problem was solved using the conditional-gradient algorithm [55], as implemented in the Python Optimal Transport (POT) library [56]. Intuitively, the optimal coupling redistributes the probability mass of the source subject’s occupancy distribution among latent states to match the target subject’s occupancy distribution, while introducing the smallest possible departure from the source subject’s reference state-transition dynamics. Higher costs indicate that matching the target occupancy distribution requires a greater modification of those dynamics. The outcome of this framework is a subject-by-subject cost matrix *C*, whose entries correspond to the transition costs *C*_*sr*_ from source subject *s* to target subject *r*, obtained as in Eq. (11). Since exchanging source and target changes the reference joint transition distribution and the marginal constraints, the resulting costs are generally asymmetric. For clustering within the MS cohort, the directional cost matrix was symmetrized by averaging reciprocal entries, i.e., 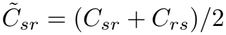.

Agglomerative clustering with average linkage was then applied to 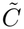. Two-, three-, and four-cluster solutions were evaluated using the silhouette score, and the solution with the highest silhouette score was selected. To assess robustness to the symmetrization choice, clustering was repeated using min(*C*_*sr*_, *C*_*rs*_) and max(*C*_*sr*_, *C*_*rs*_) as alternative symmetric dissimilarities, and agreement between partitions was quantified using the adjusted Rand index (ARI; see S1 Supporting Information). For visualization, the symmetrized dissimilarity matrix was embedded in two dimensions using multidimensional scaling (MDS), a dimensionality-reduction technique.

## Statistical analysis

All statistical tests were two-sided with a significance level of *α* = 0.05. Group differences in HMM-derived metrics between patients with MS and HCs were evaluated using Mann–Whitney *U* -tests, with Benjamini–Hochberg correction to control the false discovery rate (FDR) across all performed comparisons.

Associations between HMM-derived metrics and clinical or MRI variables within the MS cohort were assessed using Spearman rank correlations with FDR correction. Clinical and structural differences between the two OT-derived MS subgroups were assessed using Welch’s *t*-tests for SDMT and BPF, and for lesion load after log(1 + *x*) transformation to reduce its strong positive skewness. EDSS was assessed using the Mann–Whitney *U* -test, given its ordinal scale. FDR correction was applied across the four primary clinical outcomes (EDSS, BPF, lesion load, and SDMT). Covariate-adjusted analyses were additionally performed using ordinary least-squares regression with heteroskedasticity-consistent standard errors of type HC3 for continuous outcomes and proportional-odds ordinal logistic regression for EDSS. Models were adjusted for age and sex to account for potential demographic confounding, with education (years) additionally included for SDMT. FDR correction was applied across the four covariate-adjusted clinical models.

## Results

### Characterization of hidden brain states

A group-level zero-mean Gaussian HMM was fitted jointly across patients and controls, yielding *K* = 3 recurrent brain states with distinct covariance structures (Fig 2). The state covariance matrices estimated in the PCA-reduced space were projected back to the original ROI space and normalized to obtain FC matrices. A model-based mean FC was derived by weighting the state covariances according to their overall posterior occupancy and normalizing the resulting covariance matrix to correlations (Fig 2A). This estimate closely matched the empirical mean FC across participants (Pearson’s *r* = 0.996, computed after vectorizing the upper-triangular elements of the two matrices, excluding the diagonal). To characterize state-specific differences from this average organization, the model-based mean FC was subtracted from each state-specific FC matrix, yielding ΔFC matrices representing relative increases and decreases in connectivity (Fig 2C).

**Fig 2.**
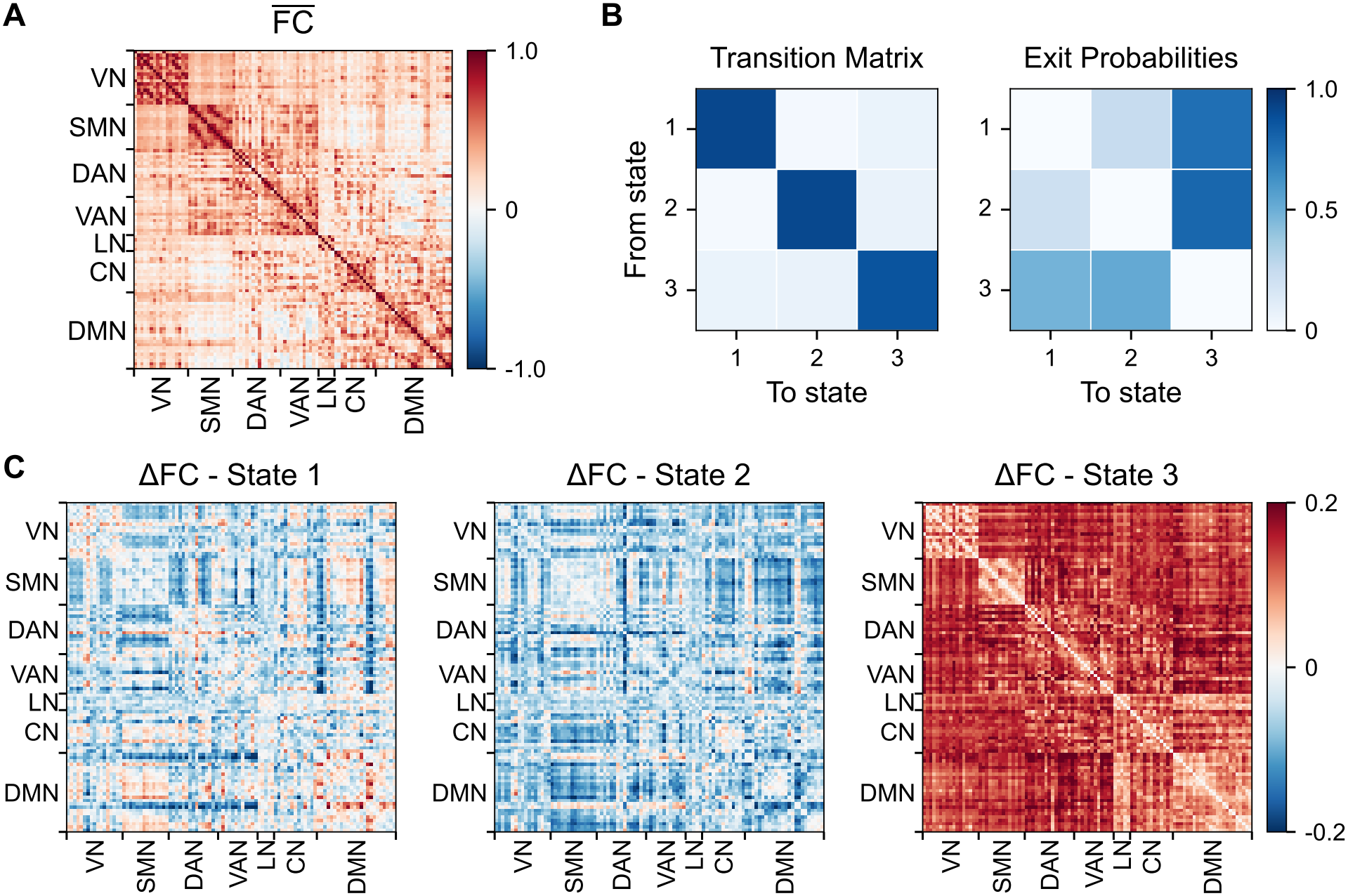
HMM-derived brain states and transition dynamics. (A) *HMM-reconstructed functional connectivity (FC)*. Average FC reconstructed from the group-level HMM in the original ROI space, obtained by combining the state-specific covariance matrices according to their overall posterior occupancy and normalizing the resulting covariance matrix to correlations. (B) *State-transition dynamics*. HMM transition matrix showing the probabilities of transitions between the three latent states. Exit probabilities were obtained by excluding self-transitions and renormalizing the remaining transition probabilities from each state, thus representing the probability of transitioning to each other state conditional on a state transition occurring. (C) *State-specific FC differences (*ΔFC). Differences between state-specific FC matrices, reconstructed in the original ROI space from the HMM covariance matrices (Σ_*k*_) and normalized to correlations, and the HMM-reconstructed average FC in (A). Positive values indicate stronger connectivity relative to the average FC, while negative values indicate weaker coupling. Network labels: VN, visual; SMN, somatomotor; DAN, dorsal attention; VAN, ventral attention; LN, limbic; CN, control; DMN, default mode.

In State 1, temporal regions of the default mode network (DMN) showed stronger coupling with non-temporal DMN components, particularly parietal and posterior midline DMN regions, and weaker coupling with ventral attention, dorsal attention, and somatomotor networks (VAN, DAN, and SMN, respectively). Together, these changes indicate a relative shift of temporal DMN coupling toward other DMN components and away from attention and sensorimotor systems. SMN regions showed a related pattern, with weaker coupling to DAN, VAN, and visual network (VN), together with stronger coupling with several non-temporal DMN regions.

State 2 showed broadly lower FC than the model-based mean, with pronounced reductions in connections from temporal DMN and SMN regions to non-temporal DMN and control network (CN) regions (Fig 2C). Compared with State 1, temporal DMN regions showed weaker coupling with non-temporal DMN components and relatively stronger coupling with VAN, DAN, and SMN (Fig 2C; see also Fig E(B) in <u>S1</u> Supporting Information), reflecting a shift in their functional coupling profile from other DMN components toward attention and sensorimotor systems. SMN connectivity showed a related but less uniform pattern, with weaker coupling to non-temporal DMN and CN regions and more heterogeneous changes across attention and visual systems.

State 3 was instead characterized by widespread positive FC deviations across within- and particularly between-network connections (Fig 2C). Despite this broad increase, the state-specific FC matrix retained a clear network structure (Fig E(A) in S1 Supporting Information). This configuration was associated with markedly lower spatial variability of BOLD activity across parcels, while global signal magnitude remained close to its average level. High-DVARS and high-FD frames occurred at rates comparable to their overall baselines, providing no evidence that this state preferentially captured frames with poorer data quality (Fig F(A,B) in S1 Supporting Information). Overall, State 3 represented a recurrent high-connectivity configuration with reduced network differentiation, consistent with a more integrated large-scale functional organization.

The transition matrix showed high self-transition probabilities for all three states, consistent with their persistence across consecutive time points (Fig 2B). Transitions away from States 1 and 2 were preferentially directed toward State 3, whereas those from State 3 were distributed between the other two states. This organization suggests that State 3 may serve as an intermediate configuration between States 1 and 2.

### Group-level differences in brain-state dynamics

We compared HMM-derived metrics between patients with MS and HCs (Fig 3A). Significant group differences were limited to States 1 and 2, whereas State 3 metrics were comparable. Patients showed lower FO in State 1 and higher FO in State 2, together with longer DT in State 2 (Mann–Whitney *U* -tests, all FDR-corrected *p <* 0.05). SR did not differ significantly between groups, indicating a comparable overall frequency of transitions between states. Together, these findings indicate a shift in temporal balance from State 1 toward State 2 in MS. This pattern reflects greater expression of the characteristic coupling profile of State 2, in which, relative to State 1, temporal DMN regions are more weakly coupled to non-temporal DMN components and more strongly coupled to attention and sensorimotor systems. In contrast, no individual static FC edge showed a significant MS–HC difference after FDR correction (Fig B in S1 Supporting Information), suggesting that the dynamic-state representation captures group differences that are not apparent in static FC.

**Fig 3.**
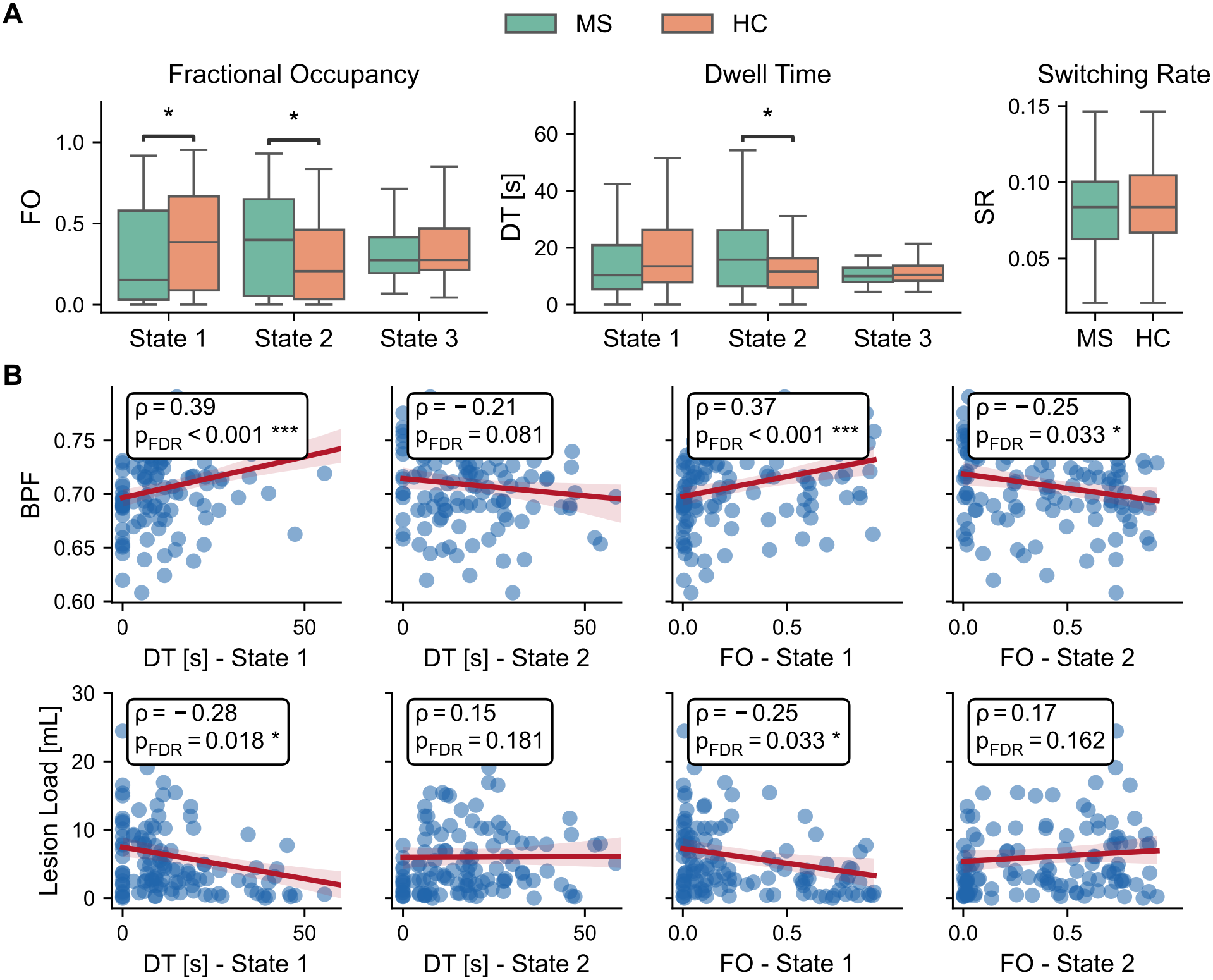
HMM-derived metrics and clinical correlations in MS. (A) *Group comparison of HMM-derived metrics*. Fractional occupancy (FO), dwell time (DT), and switching rate (SR) were compared between patients with MS and HCs. Patients showed higher FO and DT in State 2 and lower FO in State 1, while no significant group differences were observed for State 3 or SR. (B) *Clinical correlations in MS*. Spearman correlations between HMM-derived metrics and EDSS, SDMT, BPF, and lesion load were computed within the MS group, with false discovery rate (FDR) correction for multiple comparisons. Higher FO and DT in State 1 correlated positively with BPF and negatively with lesion load, whereas FO in State 2 correlated negatively with BPF. In (B), FDR-adjusted *p*-values are reported as *p*_FDR_. Statistical significance after FDR correction is denoted by * *p <* 0.05, ** *p <* 0.01, and *** *p <* 0.001.

To assess the clinical relevance of these differences in brain-state dynamics and their association with structural disease burden, we examined correlations between HMM-derived metrics and EDSS, SDMT, BPF and lesion load in patients. Among the four variables examined, only BPF and lesion load showed associations that remained significant after FDR correction; correlations with these variables are shown in Fig 3B. Higher FO and DT in State 1 were associated with higher BPF (*ρ* = 0.37, *p <* 0.001; *ρ* = 0.39, *p <* 0.001, respectively) and lower lesion load (*ρ* = *−*0.25, *p* = 0.033; *ρ* = *−*0.28, *p* = 0.018, respectively). Higher FO in State 2 was instead associated with lower BPF (*ρ* = *−*0.25, *p* = 0.033). These associations link the State 1–State 2 temporal balance to structural disease burden.

### Patient stratification in MS based on brain-state dynamics

We next used OT to quantify pairwise dissimilarities in HMM-derived brain-state dynamics within the MS cohort, yielding the matrix shown in Fig 4A. Each entry represents the cost of reaching the target subject’s latent-state occupancy distribution from that of the source subject, with larger costs indicating a greater required departure from the source subject’s state-transition dynamics. Because the OT costs are inherently directional (*C*_*sr*_ *≠C*_*rs*_), the matrix was symmetrized before clustering, as described in Materials and methods. Agglomerative clustering identified a two-cluster partition within the MS cohort, which achieved the highest silhouette score (0.58, compared with 0.46 and 0.43 for three and four clusters, respectively). For visualization, MDS was applied to the symmetrized OT matrix to obtain a two-dimensional representation of the pairwise relationships among subjects. The resulting MDS dimensions (MDS1 and MDS2) were strongly associated with HMM-derived FO values (Fig 4B; Fig C in S1 Supporting Information), consistent with the low-dimensional representation of the OT dissimilarity structure reflecting inter-individual variation in latent brain-state dynamics. Alternative symmetrizations based on min(*C*_*sr*_, *C*_*rs*_) and max(*C*_*sr*_, *C*_*rs*_) produced identical subject assignments (ARI = 1.0; S1 Supporting Information), indicating that the partition was robust to the specific symmetrization choice. Cluster assignments were also highly consistent across the tested HMM configurations (*M* = 30, 40; *K* = 3, …, 6), with pairwise ARI values ranging from 0.77 to 1.00 (Fig G in S1 Supporting Information), suggesting that the stratification was stable across these model specifications.

**Fig 4.**
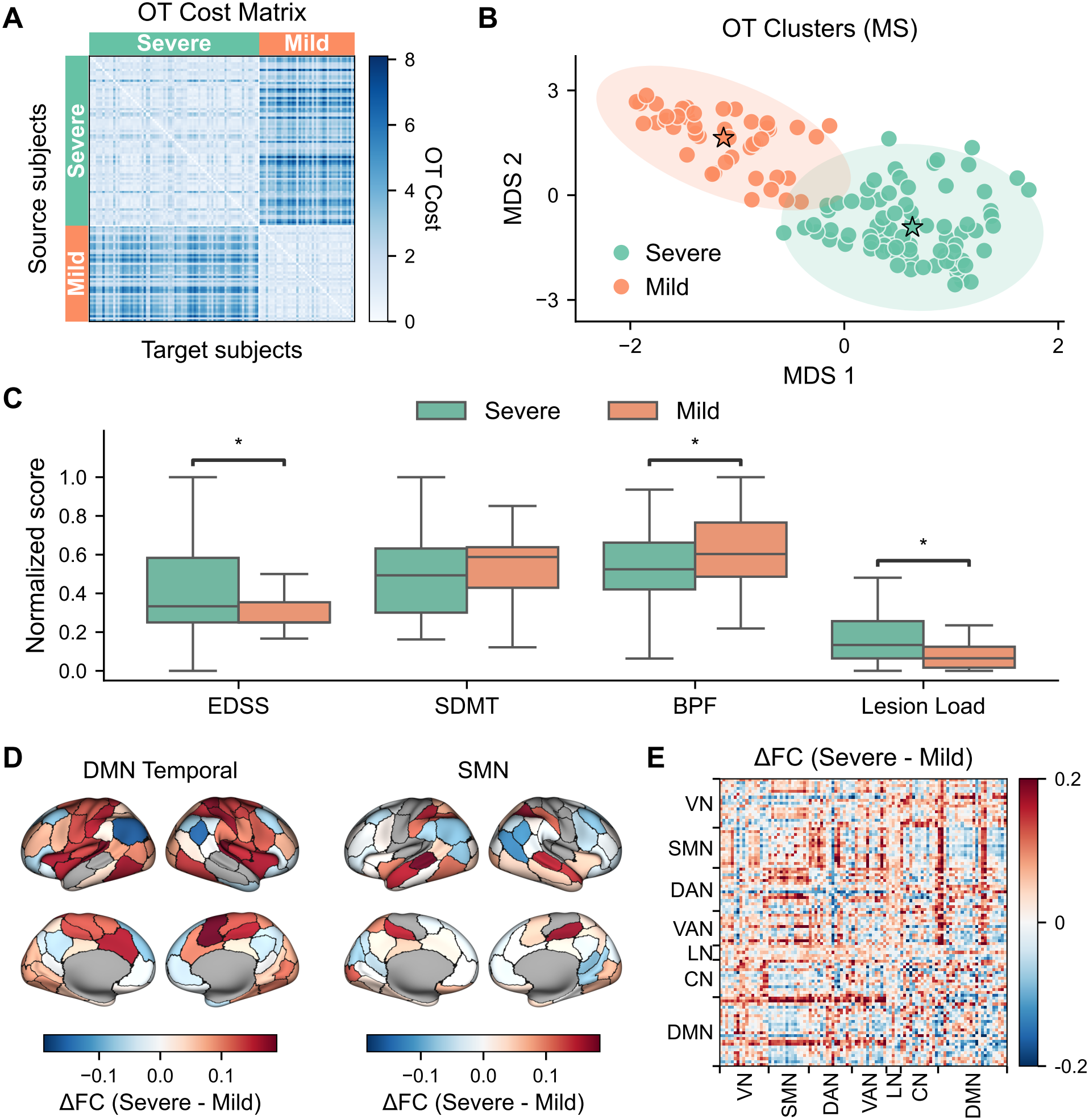
Data-driven patient stratification in MS based on brain-state transition costs. (A) *OT cost matrix*. Pairwise directional transition costs derived from the Schrödinger bridge framework, based on HMM-derived brain dynamics. (B) *OT-derived MS clusters*. Agglomerative clustering applied to the symmetrized OT cost matrix identifies two patient groups, visualized in a low-dimensional space using multidimensional scaling (MDS). Stars indicate cluster centroids, and shaded ellipses represent 95% Gaussian covariance contours. (C) *Clinical and structural group comparison*. Compared to the *Mild* group, the *Severe* group shows higher EDSS, lower BPF, and greater lesion load, while no significant difference is observed in SDMT. Statistical significance after FDR correction is denoted by * *p <* 0.05. For visualization, clinical and structural measures were min-max normalized. (D) *Network-specific connectivity profiles*. Mean FC differences between the *Severe* and *Mild* groups are shown for bilateral temporal DMN and SMN parcels (shown in gray). For each network, FC differences between the corresponding bilateral parcels and the rest of the cortex were averaged across those parcels and mapped onto the cortical surface. (E) *Whole-brain FC differences*. Difference between the mean FC matrices of the *Severe* and *Mild* groups (ΔFC, *Severe*–*Mild*). Positive values indicate stronger connectivity in the *Severe* group. Cluster labels (*Mild* and *Severe*) used throughout the figure were assigned *a posteriori* based on the differences observed in (C).

The two clusters differed in clinical and MRI-derived structural measures (Fig 4C) and were therefore labeled *Mild* and *Severe* a posteriori. Compared with the *Mild* group, the *Severe* group had higher EDSS scores (rank-biserial *r* = 0.23, *p* = 0.039), lower BPF (Hedges’ *g* = *−*0.47, *p* = 0.022), and greater lesion load (Hedges’ *g* = 0.54, *p* = 0.022), whereas SDMT did not differ significantly between groups (Hedges’ *g* = *−*0.25, *p* = 0.191). Differences in EDSS, BPF, and lesion load remained significant after adjustment for age and sex (all *p* = 0.041), whereas the SDMT difference remained non-significant after additional adjustment for education (*p* = 0.437). All reported *p*-values were FDR-adjusted within the corresponding set of four tests. Importantly, clustering based on static FC embeddings obtained with PCA or Uniform Manifold Approximation and Projection did not recover subgroups with comparable clinical and structural differentiation (Fig D in S1 Supporting Information), suggesting that the OT-based representation of latent brain-state dynamics captured information not identified by static FC-based stratification.

To further characterize the functional organization of the two OT-derived groups, we examined differences between their group-average static FC matrices (Fig 4D,E). The most prominent network-specific difference involved temporal DMN regions, which in the *Severe* group showed weaker coupling particularly with parietal and posterior midline DMN regions, but stronger coupling with attention and sensorimotor systems.

SMN regions also showed stronger coupling with posterior parietal DAN regions and medial VAN regions in the midcingulate cortex. At the whole-brain level, the difference between the group-average FC matrices (*Severe*–*Mild*) closely resembled the difference between the State 2 and State 1 FC matrices (Pearson’s *r* = 0.817; Fig E(B) in S1 Supporting Information). Overall, these findings link the MS-associated shift in the temporal balance between States 1 and 2 to clinical heterogeneity, with the principal FC reorganization separating these states also expressed across the clinical *Mild* –*Severe* axis.

## Discussion

In this study, we developed a novel framework to capture inter-individual variability in resting-state brain dynamics and applied it to derive a data-driven stratification within RRMS. The dynamical modeling was based on an HMM, which jointly estimates recurrent latent brain states characterized by distinct connectivity patterns and the transition probabilities governing their temporal organization [18, 19, 44]. This probabilistic approach avoids the need to select a predefined window length and to identify recurrent connectivity states through a separate clustering step that does not account for the temporal ordering of the windowed FC patterns, thereby addressing well-recognized limitations of sliding-window dynamic functional connectivity (dFC) methods [14, 21]. Moreover, because the same set of states is estimated across participants, individual differences in state occupancy and transition dynamics can be characterized within a common latent-state space [18]. Accordingly, HMM studies have typically relied on dynamic metrics such as fractional occupancy, dwell time, and switching rate for group comparisons or prediction.

Here, we build on this feature-based description by combining the HMM-derived representation with a Schrödinger bridge formulation of OT to compare latent-state dynamics across participants. Within this framework, we quantify the minimal modulation of one participant’s stochastic transition dynamics required to reproduce the latent-state occupancy distribution of another’s, yielding a pairwise dissimilarity measure grounded in the probabilistic representation provided by the HMM. This formulation explicitly accounts for the inherent stochasticity of neural dynamics, which is important for accurately estimating brain-state transition costs [26]. This contrasts with network-control approaches based on deterministic dynamical systems, in which control cost is defined as the minimum input energy required to drive the system between specified brain states [24, 57]. In MS, Broeders et al. [58] recently applied this perspective, using subject-specific structural networks to quantify the control energy required for transitions between functional connectivity states, and found higher values in cognitively impaired patients. Whereas such approaches characterize transitions between states *within* individual participants, our formulation defines a transition cost *between* participants, and the resulting dissimilarity matrix forms the basis for the unsupervised stratification of the MS cohort. In a supplementary analysis, clustering directly on HMM-derived occupancy and transition measures recovered the same partition (Fig H in S1 Supporting Information), indicating that the identified group structure is already encoded in the underlying HMM representation. Nonetheless, OT provides a theoretically motivated geometry for comparing inter-individual differences, and its stochastic-control interpretation places the framework within broader views of brain-state transitions in terms of energetic or probabilistic costs [24, 25]. More broadly, the present approach provides a tool for subject-level dynamic phenotyping, contributing to the emerging landscape of dynamic approaches for capturing the complex nature of behavior and brain disorders [59].

Application of this framework to RRMS revealed an altered temporal organization of recurrent brain states relative to healthy controls, centered on a shift in the balance between States 1 and 2, while no significant group difference in switching rate was detected. The connectivity patterns distinguishing these states reflect a selective reorganization of functional relationships across transmodal/default-mode systems and externally oriented attention and sensorimotor networks. Temporal DMN regions provide the most coherent subsystem-level expression of this reorganization, showing weaker coupling with other DMN components and stronger coupling with attention and sensorimotor systems in State 2 compared with State 1, alongside stronger SMN coupling with posterior parietal DAN and midcingulate VAN regions. Although these changes involve specific network components, they are consistent with a broader alteration of sensory-to-transmodal functional organization, as suggested by the compression of the corresponding functional gradient reported in a complementary analysis of the same cohort [35]. The involvement of specific DMN components is consistent with the organization of this network into functionally distinct subsystems [60] and with previous evidence that connectivity among DMN subregions is altered in RRMS [61–63], including marked alterations in the temporal pole [62]. More broadly, recent meta-analytic evidence identified both increased and decreased FC in distinct regions of the DMN and SMN in MS, underscoring the spatial heterogeneity of functional alterations within these networks [64]. In the present cohort, the temporal expression of the State 1–State 2 balance covaried with BPF and lesion load, linking this altered dynamical organization to structural disease burden. These findings add to a growing dFC literature in MS relating brain network dynamics to cognitive impairment, disability, and disease phenotype. Reported relationships broadly concern two aspects of brain dynamics: alterations in the temporal expression and connectivity of recurrent functional configurations [36, 65, 66], and changes in the stability and reconfiguration of functional networks over time, whether assessed globally [37] or at the level of individual networks and regions [67–69]. Although the specific abnormalities differ across dynamical measures and patient subgroups, these studies collectively indicate that time-varying functional organization represents a clinically relevant dimension of functional alteration in MS.

At the inter-individual level, this clinical relevance was reflected in the OT-derived stratification within the MS cohort. The two identified groups differed significantly in disability, BPF, and lesion load despite sharing the same relapsing-remitting MS diagnosis, supporting the biological relevance of this dynamical stratification. Crucially, clustering of linear and nonlinear static-FC embeddings did not yield groups with comparable clinical or structural differentiation, indicating that temporal dynamics captured inter-individual heterogeneity that was not recovered by the static representations tested here. This finding is consistent with previous MS studies showing that dFC measures can better distinguish disability status and reveal clinically relevant associations not captured by time-averaged approaches [38, 65]. Importantly, the dynamical stratification was also reflected in group-averaged static FC, with the FC difference between the *Severe* and *Mild* groups closely resembling the State 2–State 1 pattern (*r* = 0.817). This strong correspondence indicates that the major connectivity organization distinguishing the recurrent states was also expressed across the *Mild* –*Severe* axis, consistent with inter-individual differences in state expression being reflected in average connectivity. These findings extend previous data-driven MRI phenotyping efforts in MS, which have predominantly relied on structural MRI features [31–33, 70], by showing that functional brain dynamics can define an additional axis of patient heterogeneity.

Several limitations should be considered when interpreting the present findings and in view of future developments of the framework. First, the relatively short duration of the resting-state acquisitions (approximately six minutes) constrains the precision of subject-specific estimates of state occupancy and transition dynamics, and consequently the reliability of the resulting pairwise dissimilarities. Longer acquisitions would provide more reliable state-based estimates, while repeated scans would allow their consistency across sessions to be assessed directly [71]. Although such protocols are often challenging to implement in clinical settings, they provide an important benchmark for dynamic phenotyping, given evidence for reproducible individual differences in brain-state dynamics across repeated resting-state measurements [72] and the need to establish temporal stability and identifiability of model-derived measures used as phenotypes [73]. Second, although resting-state fMRI allows spontaneous brain dynamics to be characterized without constraining the analysis to a predefined cognitive domain, the absence of an experimentally controlled cognitive context limits the functional interpretation of the inferred recurrent configurations. Their temporal expression cannot be directly linked to specific cognitive processes, and the potential contribution of transient variations in ongoing cognition to disease-related differences cannot be fully disentangled. Finally, the cross-sectional, single-cohort design limits the generalizability of the inferred dynamical organization and OT-derived stratification. Independent longitudinal and multicenter studies will be required to assess their evolution with disease progression, reproducibility across cohorts, and potential modulation by treatment.

## Conclusion

In summary, this study shows that combining HMM-based modeling of recurrent brain states with a Schrödinger bridge formulation of OT provides a pairwise measure of dissimilarity in latent-state dynamics that captures clinically and structurally relevant heterogeneity in RRMS. While several methodological and practical challenges remain, particularly concerning scan duration, test–retest reliability, and generalizability, our results establish the feasibility and clinical relevance of transport-theoretic dynamic phenotyping. If validated in independent longitudinal cohorts, such dynamical signatures could be evaluated for prognostic stratification and treatment-response prediction. More generally, the framework could be extended to other neurological and psychiatric conditions characterized by heterogeneous clinical presentations. As neuroimaging datasets continue to grow in size and temporal resolution, transport-based approaches to brain dynamics may provide increasingly refined descriptions of individual differences in brain function and their relationship to clinical phenotypes.

## Declarations

### Author contributions

Conceptualization: LT, GB, BM, FC, SDM, AV, MAll, AB, SS, MZ. Methodology: LT, GB, BM, FC, MAll, AB, SS, MZ. Investigation: LT, GB, BM, FC. Visualization: LT, GB, FC. Supervision: FE, SDM, AV, MAll, AB, SS, MZ. Data curation: AG, MAlt, APDR, AD, FE. Writing – original draft: LT, GB, FC, MAll, MZ. Writing – review and editing: LT, GB, BM, FC, AG, MAlt, APDR, AD, FE, SDM, AV, MAll, AB, SS, MZ.

### Data and code availability

The clinical and neuroimaging data analyzed in this study were obtained from the cohort previously described by De Rosa et al. [35]. In compliance with Italian data protection regulations governing patient privacy, the raw data cannot be made publicly available. However, anonymized data can be made available to qualified researchers upon reasonable request to the corresponding author of the original study (FE).

Access may require a formal data-sharing agreement.

All code developed for this study is publicly available at https://github.com/LTaffarello/hmm-ot.

### Ethics statement

As detailed in [35], ethical approval was received from the local ethical standards committee, and written informed consent was obtained from all participants after the nature and possible consequences of the study were fully explained.

## Supporting information

**S1 Supporting Information. Supplementary methods, analyses, and figures**. Further details on HMM model selection and brain-state characterization, quality-control assessments, robustness of OT-based clustering, and comparisons with clustering approaches based on static FC and HMM-derived features.

