## Supplementary material for "Discovery of latent clinical phenotypes through optimal transport of brain dynamics": S1 Supporting Information

Taffarello et al.

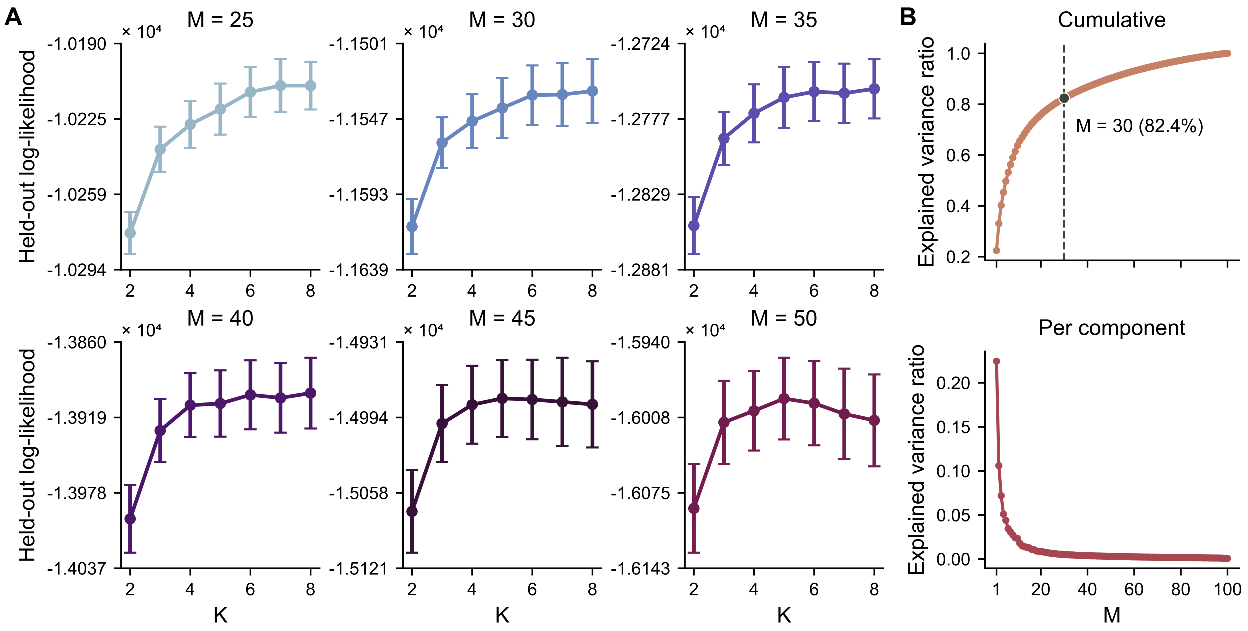


**Fig A. HMM model selection and PCA dimensionality.** (A) *HMM model selection.* Mean held-out marginal log-likelihood per subject across the five folds of subject-wise cross-validation, shown as a function of the number of HMM states ($K = 2, \ldots, 8$) for different PCA dimensionalities ($M = 25, 30, 35, 40, 45, 50$). Error bars indicate the standard error across folds. Across PCA dimensionalities, the largest increase in held-out log-likelihood occurred between $K = 2$ and $K = 3$, whereas additional states yielded only modest gains. (B) *PCA explained variance.* Explained variance ratio for individual principal components and cumulative explained variance as a function of the number of components. The $M = 30$ components retained for the primary analysis explained 82.4% of the total variance.

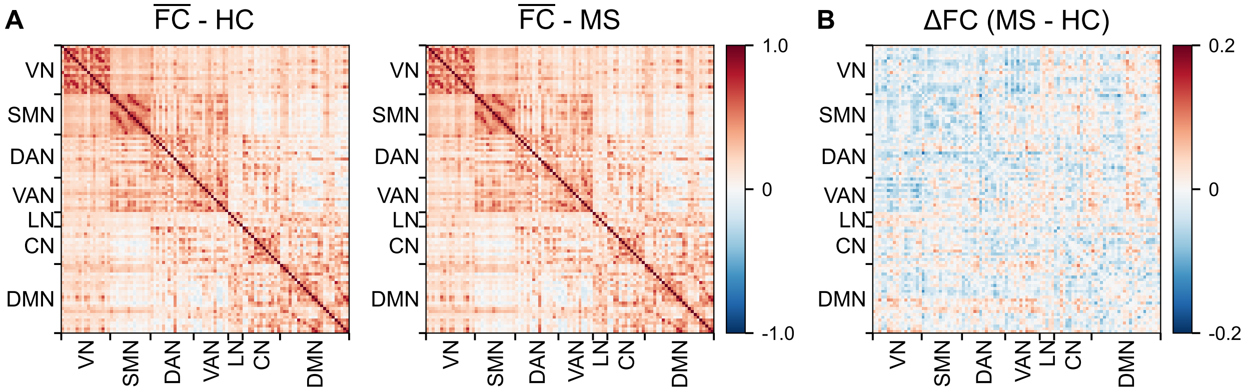


**Fig B. Group-average static FC in MS and HC.** (A) *Group-average FC.* Mean static FC matrices for HC and MS, obtained by averaging subject-level FC matrices within each group. (B) *MS-HC FC differences.* Difference between the group-average FC matrices ($\Delta$FC, MS $-$ HC). Positive values indicate stronger mean connectivity in MS, whereas negative values indicate stronger mean connectivity in HC. For statistical inference, individual FC edges were Fisher-transformed and compared between groups using edge-wise Welch's $t$-tests with Benjamini-Hochberg FDR correction. No individual edge was significant after FDR correction ($p < 0.05$).

### Robustness of OT-based clustering to matrix symmetrization

Because Schrödinger bridge OT costs are directional, the resulting subject-by-subject cost matrix is asymmetric. To quantify the degree of asymmetry, we computed the symmetry degree

$$1 - \frac{\|C - C^{\top}\|_F}{\|C\|_F}, \tag{S1}$$

where $\|\cdot\|_F$ denotes the Frobenius norm. In the MS cohort, the symmetry degree was 0.745, indicating that the cost matrix remained largely symmetric despite the directionality of the OT costs. Before clustering, we derived a symmetric dissimilarity matrix averaging reciprocal entries $\tilde{C}_{sr} = (C_{sr} + C_{rs})/2$. To assess whether the clustering outcome depended on the specific symmetrization strategy, we repeated the agglomerative clustering using two alternative symmetric constructions, namely $\min(C_{sr}, C_{rs})$ (best-case) and $\max(C_{sr}, C_{rs})$ (worst-case) reciprocal dissimilarities. Across all three approaches (mean, min, max), silhouette analysis consistently selected a two-cluster solution (silhouette scores ranging from 0.57 to 0.60), and subject assignments were identical (ARI = 1.0). Despite this identical partitioning, local geometric properties differed across symmetrizations. Specifically, the maximum change in pairwise dissimilarities reached 35% relative to the mean symmetrization, while 83% (mean vs. max) and 74% (mean vs. min) of subjects retained the same sets of five nearest neighbors, indicating some local rearrangements of neighborhood structure. In contrast, the global geometry of the distance matrices was highly preserved, with correlations between upper-triangular entries exceeding 0.98 for mean vs. max and mean vs. min (and 0.96 for max vs. min). Because average-linkage agglomerative clustering relies on average inter-cluster dissimilarities rather than local neighborhood relations, the preserved global structure is consistent with the identical final partitions obtained across symmetrization schemes. Together, these analyses demonstrate that the identified MS subgroups are driven by stable large-scale organization in OT-derived dissimilarities and are robust to the specific choice of symmetrization, despite modest directional asymmetry and local geometric variations.

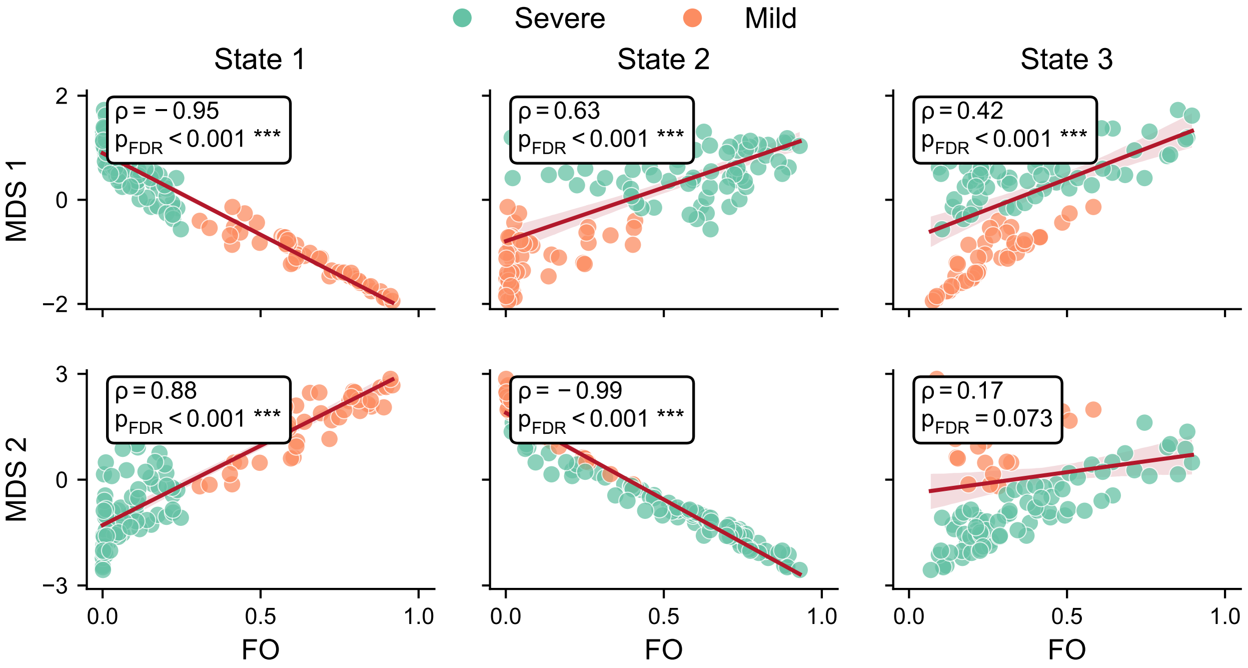


**Fig C. Associations between OT-derived MDS dimensions and FO in MS.** Spearman correlations between the first two MDS dimensions (MDS1 and MDS2), obtained from the OT-based subject dissimilarity matrix, and FO of each HMM state within the MS group. MDS1 was strongly associated with FO across all three states, whereas MDS2 showed strong associations with FO in States 1 and 2 but not in State 3. Subjects are color-coded based on their cluster assignment (*Mild* vs *Severe*). Reported $p$-values were FDR-corrected across all tested correlations between MDS dimensions and HMM-derived metrics. Statistical significance after FDR correction is denoted by * $p < 0.05$, ** $p < 0.01$, and *** $p < 0.001$.

### FC-based clustering

As a comparison to the OT-based stratification, we repeated the clustering analysis using static FC. For each patient with MS, FC matrices were vectorized and assembled into a subject-by-feature matrix, followed by dimensionality reduction using PCA. Ward-linkage agglomerative clustering was then applied to the PCA representations, and a two-cluster solution was selected for consistency with the OT-based analysis and guided by silhouette scores. The resulting PCA-derived clusters were compared in terms of clinical and structural measures, and no significant group differences were observed. Comparable results were obtained across several PCA dimensionalities (2, 5, 10, 25, 60, and 121 components), suggesting that the limited clinical separation was not driven by a specific choice of embedding dimension. We further evaluated a nonlinear embedding of static FC features using Uniform Manifold Approximation and Projection (UMAP), followed by Ward-linkage agglomerative clustering into two groups. Because UMAP embeddings can vary across stochastic runs, we used a consensus strategy based on 500 repetitions of the embedding and subsequent clustering. The resulting co-association matrix was converted to a dissimilarity matrix and clustered using average linkage to obtain the final consensus partition. Also in this case, group comparisons did not reveal significant differences in clinical or structural measures. Repeating the consensus analysis using 2-, 5-, and 10-dimensional UMAP embeddings yielded the same conclusion. Overall, these analyses suggest that static FC-based clustering provides weaker clinical separation than OT-based stratification that incorporates the temporal organization of brain-state dynamics.

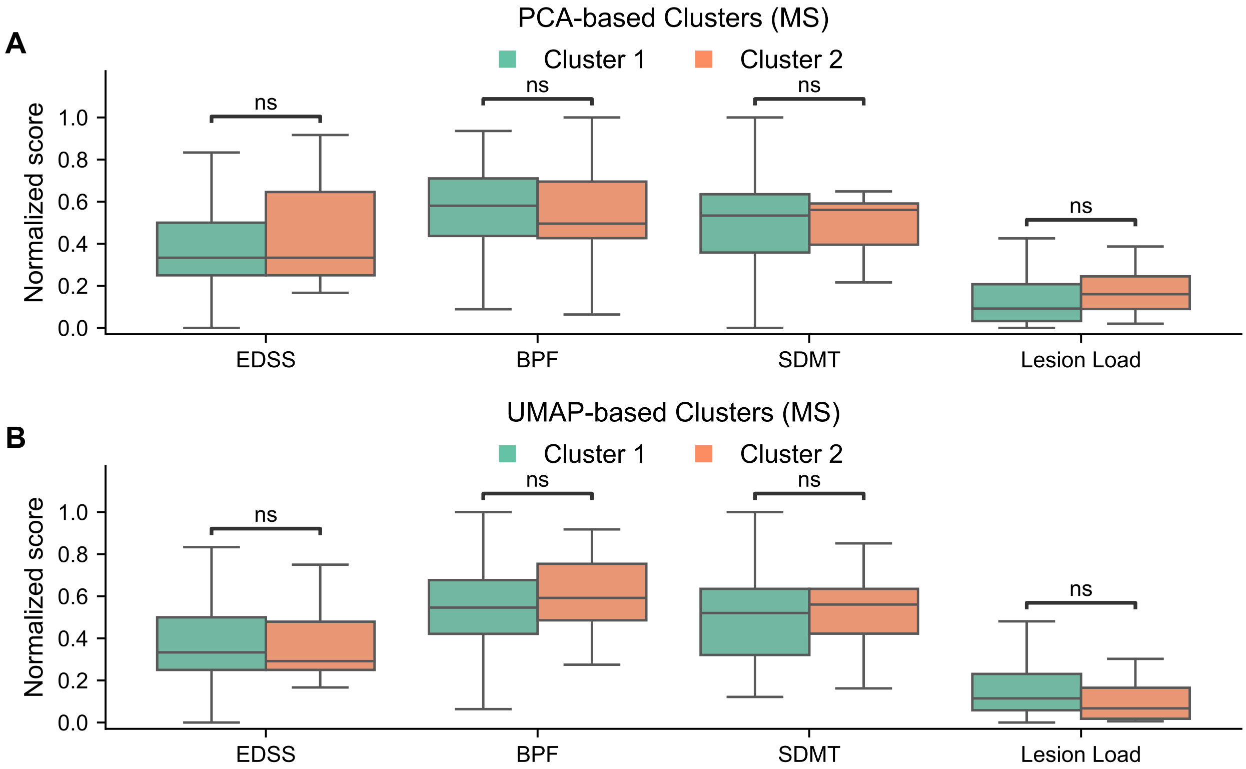


**Fig D. Static FC-based clustering within the MS cohort.** Boxplots summarize clinical and structural measures across clusters obtained by two-group agglomerative clustering based on PCA (5 PCs; A) and UMAP-based consensus clustering (5 dimensions; B) of static FC features in patients with MS. Group comparisons were performed without covariate adjustment; none were significant after FDR correction (ns, not significant).

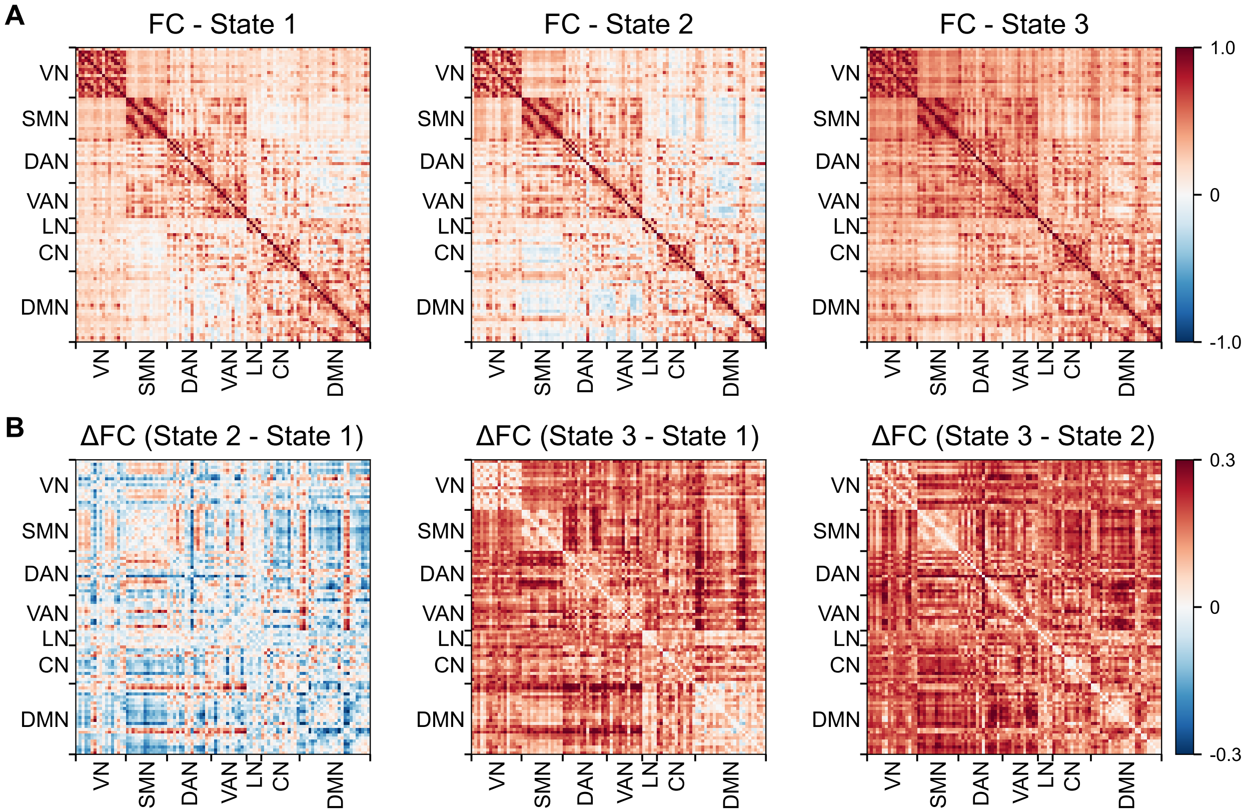


**Fig E. FC patterns across HMM states.** (A) *State-specific FC.* FC matrices reconstructed in the original ROI space from the HMM covariance matrices ($\Sigma_k$) and normalized to correlations. (B) *Between-state differences.* Pairwise differences between the reconstructed FC matrices, shown as $\Delta$FC (State 2 − State 1), $\Delta$FC (State 3 − State 1), and $\Delta$FC (State 3 − State 2).

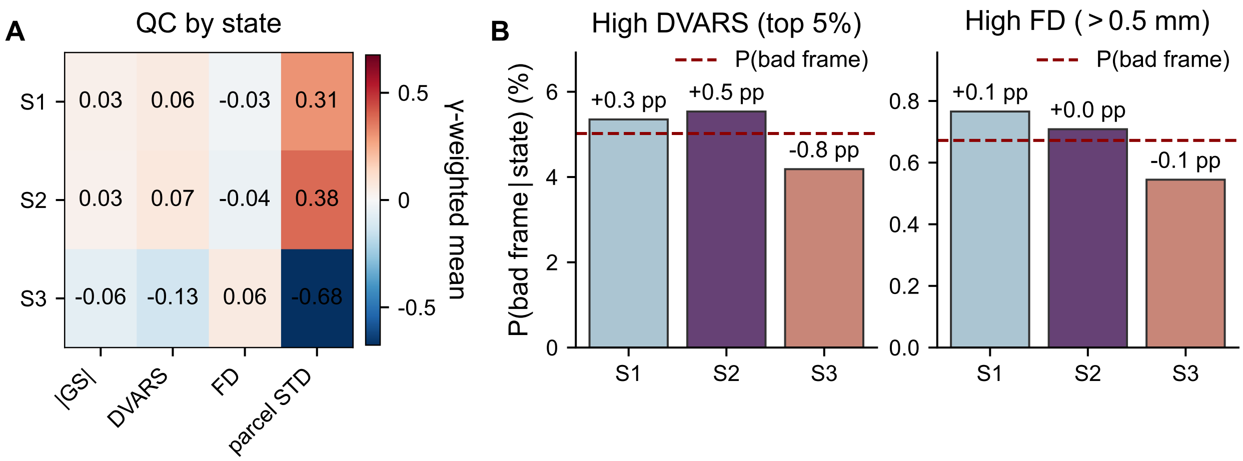


**Fig F. Quality-control assessment of HMM states.** (A) *Signal and quality-control measures by state.* Global signal (GS) magnitude ($|GS|$), computed as the absolute value of the GS at each time point, DVARS, FD, and spatial standard deviation of BOLD activity across parcels (parcel STD) were standardized within each subject and averaged across time points using the posterior state probabilities ($\gamma$) as weights. GS magnitude, DVARS, and FD showed only small differences across states, whereas parcel STD showed a stronger state dependence, consistent with the covariance structure distinguishing the HMM states. (B) *High-DVARS and high-FD frames by state.* State-conditional probabilities of high-DVARS (top 5% of DVARS values within each subject) and high-FD (FD $> 0.5$ mm) frames. Bars show the posterior-probability-weighted fraction of high-DVARS or high-FD frames within each state. Dashed lines indicate the corresponding overall probability irrespective of state, and annotations report the difference from this baseline in percentage points (pp). State-specific probabilities remained close to the corresponding overall baselines, indicating only modest differences in the occurrence of high-DVARS and high-FD frames across states.

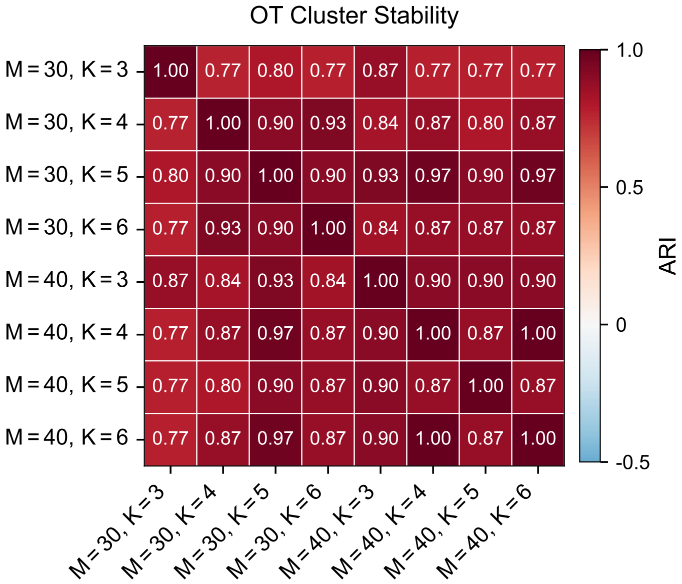


**Fig G. Robustness of OT-derived clustering across HMM configurations.** Pairwise similarity of the OT-derived MS cluster assignments across different PCA dimensionalities ($M = 30, 40$) and numbers of HMM states ($K = 3, \ldots, 6$), quantified using ARI. Cluster assignments were highly consistent across configurations, with pairwise ARI values ranging from 0.77 to 1.00, indicating that the identified patient stratification was robust to variations in PCA dimensionality and HMM model order.

### HMM feature-based clustering

To assess whether the OT-derived stratification could be recovered directly from HMM-derived features, we performed an additional feature-based clustering analysis within the MS group. For each subject, FO values and the elements of the joint transition probability matrix ($Q$) were combined into a single feature vector, and each feature was standardized across subjects. For the three-state HMM, this vector contains 12 features (three FO values and nine elements of $Q$), although these quantities are not independent: fractional occupancies sum to one, $Q$ is normalized such that $\sum_{ij} Q_{ij} = 1$, and its row and column marginals closely approximate the corresponding fractional occupancies. These constraints substantially reduce the effective dimensionality of the representation, with the first six principal components explaining 99.95% of the total variance. Ward-linkage agglomerative clustering was therefore applied to these six principal components, using a two-cluster solution for direct comparison with the OT-derived stratification. The resulting feature-based clustering recovered the same partition of patients as the OT-based clustering.

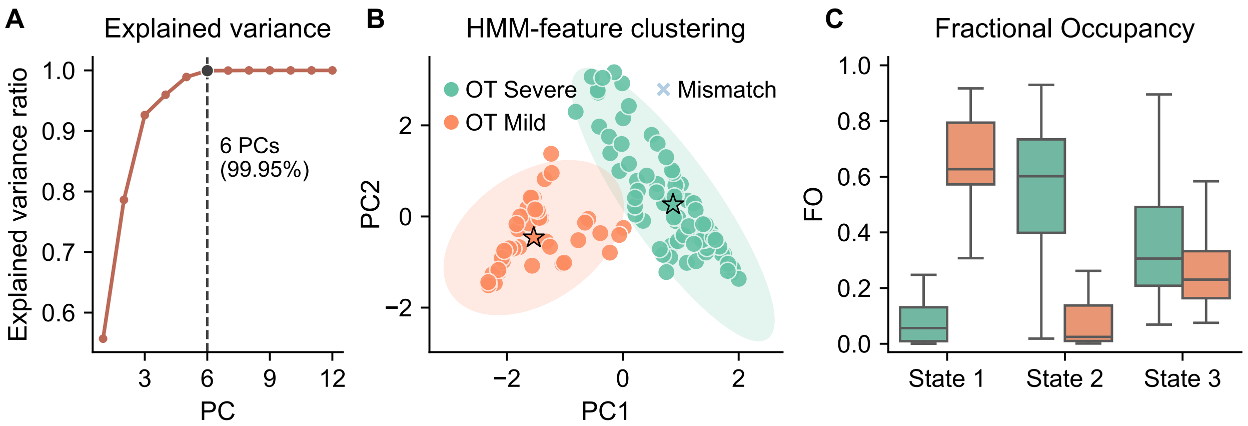


**Fig H. Relationship between HMM-derived features and OT-based patient stratification.** (A) *Feature dimensionality.* Cumulative explained variance from PCA applied to the standardized HMM-derived features, comprising FO and the elements of the subject-specific joint transition probability matrix ($Q$). The first six principal components explained 99.95% of the total variance. (B) *HMM feature-based clustering.* Agglomerative clustering of patients based on the first six principal components of the FO and $Q$ features, with subjects projected onto the first two components for visualization. Colors indicate the *Severe* and *Mild* groups defined by the OT-derived stratification. The resulting partition matched these groups. (C) *FO across patient groups.* Distribution of FO for each HMM state in the OT-derived *Severe* and *Mild* groups. Occupancy profiles differed particularly for States 1 and 2, with the *Severe* group showing lower FO in State 1 and higher FO in State 2 than the *Mild* group.

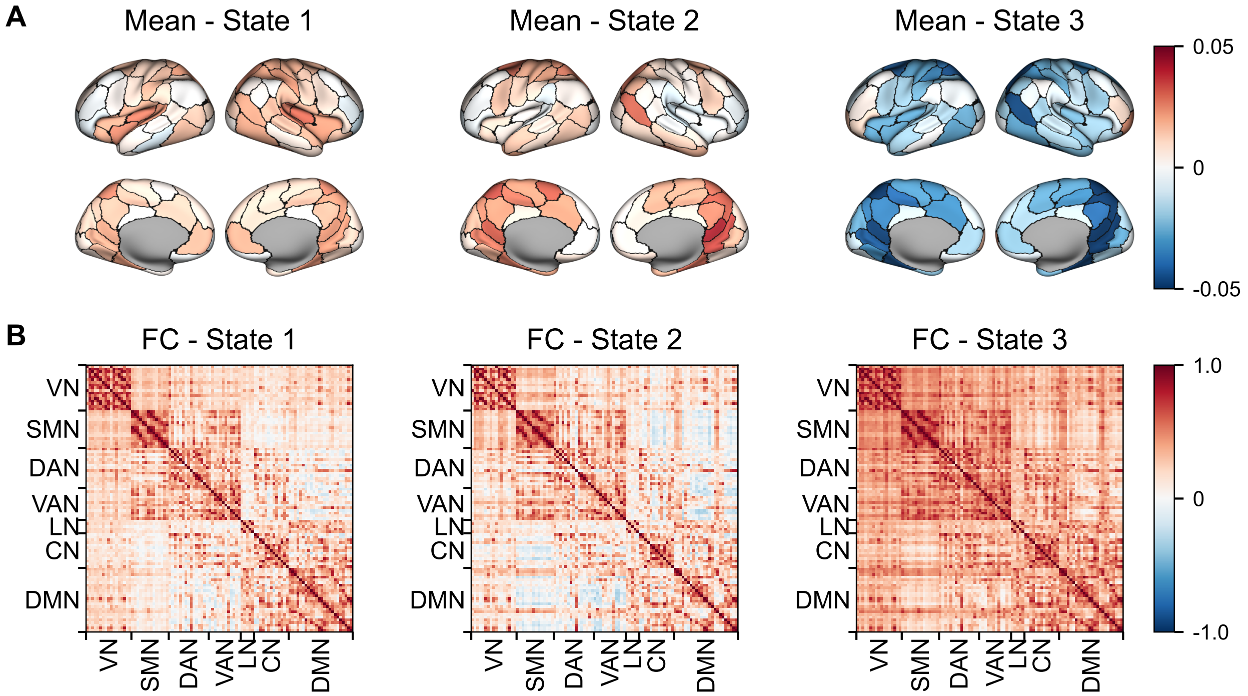


**Fig I. HMM states obtained with unconstrained state means.** Results of a three-state Gaussian HMM in which both the mean activation vectors ($\mu_k$) and covariance matrices ($\Sigma_k$) were estimated. (A) *Mean activity.* Mean activation vectors ($\mu_k$) for each state, reconstructed in the original ROI space and mapped onto the cortical surface. Their small magnitude supports the use of the zero-mean constraint in the primary HMM. (B) *FC patterns.* Matrices reconstructed in the original ROI space from the estimated covariance matrices ($\Sigma_k$) and normalized to correlations. State labels were aligned to the corresponding states of the primary zero-mean HMM for comparison.
